# Structural insights into FOXP2 multimerization and interactions from AlphaFold3 modeling

**DOI:** 10.1101/2025.04.25.650520

**Authors:** Rafal Adnan, Ali Al-Fatlawi

**Affiliations:** Biotechnology Center (BIOTEC), Center for Molecular and Cellular Bioengineering, Technische Universität Dresden, Dresden, Germany; University of Kufa, Iraq

## Abstract

FOXP2 is a transcription factor critical for speech and language, yet its full-length structure remains unresolved experimentally. We used AlphaFold3 to model full-length human FOXP2 and its complexes. AlphaFold3 predicts that FOXP2 forms a symmetric homo-hexamer, in which oligomerization orders regions that are disordered in the monomer. This assembly may facilitate cooperative DNA binding via the forkhead DNA-binding domains (FHDs). The FHD shows variable sequence affinities; high-affinity sites yield broader interfaces and greater stabilization, consistent with SPR data. Mapping of disease variants and human-specific substitutions revealed clustering in regions predicted to be stabilized by hexamer assembly. Modeling the R553H mutation showed loss of DNA contact at a critical recognition helix, which may explain its impact on transcription. We modeled FOXP2 interactions with 12 partners using AlphaFold3-multimer, evaluating interface consistency across replicates. DNA-dependent interactions were predicted with FOXP1, FOXP4, Pin1, and CDK3, involving the LZ and zinc finger domains. Other predicted interactions were low-confidence and sensitive to FOXP2’s dynamic behavior. This work generates a testable, in silico structure-function map of FOXP2 that provides a framework for future experimental validation.

## Introduction

FOXP2 belongs to the forkhead-box (FOX) family of transcription factors with key roles in neurodevelopment and human speech and language [1,2]. Dysregulated FOXP2 activity is also linked to oncogenesis, as seen in other developmental regulators like DLX5/6 [3]. However, the regulation of FOXP2’s transcriptional activity remains poorly understood [4]. FOXP2 is highly conserved; only two amino acid substitutions distinguish the human protein from that of great apes [1]. Initial reports suggested these substitutions underwent positive selection in recent human evolution [2], though more recent analyses found they arose before modern humans [5]. Notably, FOXP2 is the first gene identified to be directly involved in a human speech and language disorder: a point mutation (R553H) in FOXP2 co-segregates with speech apraxia and grammatical impairments in a large family (the KE family), and a chromosomal translocation disrupting FOXP2 causes a similar disorder in an unrelated individual [6]. Mutations in the transcription factor FOXP2 are known to cause a rare, monogenic form of severe speech and language disorder, characterized by deficits in articulation, grammar, and orofacial motor control [7]. Despite its importance, understanding FOXP2’s structure-function relationships has been challenging.

FOXP2 is a 715-amino-acid multi-domain protein with regions of high sequence conservation as well as low-complexity stretches. It contains an N-terminal polyglutamine tract, a zinc-finger motif, a leucine-zipper (LZ) domain, and a C-terminal forkhead (winged-helix) DNA-binding domain (FHD) [8]. The isolated FHD of FOXP2 has been structurally characterized in complex with DNA [8], revealing how its recognition helix inserts into the DNA major groove to contact specific base pairs [8]. FOXP proteins form both homo- and hetero-oligomers, influencing transcriptional regulation via their LZ motif [9]. The FOXP LZ is a parallel coiled-coil essential for dimerization and higher-order multimerization [10]. Biochemical studies have shown that FOXP1 and FOXP2 fragments containing the LZ form oligomers in solution (in contrast to the isolated FHD, which remains monomeric) [9]. FOXP2 in particular shows a strong propensity for higher-order oligomers - experimentally, it can assemble into complexes as large as hexamers or octamers, far beyond the basic dimer [9]. This oligomerization, while mediated by the LZ, is further modulated by intrinsically disordered regions (IDRs) of FOXP2. Indeed, >50% of FOXP2’s sequence is predicted to be disordered, and these disordered regions contribute actively to folding and multimerization [9]. Notably, FOXP2 (and FOXP1/4) has a ∼70-residue acidic tail at the extreme C-terminus that is predicted to be disordered yet is functionally important [9,10]. This acid-rich tail can fold back onto the FHD, making specific intramolecular contacts (for example, between acidic residues in the tail and basic Arg residues in the FHD) [9]. Such intramolecular interactions are thought to auto-inhibit DNA binding or oligomerization; differences between FOXP2 and FOXP1 tails correlate with differences in DNA-binding affinity and target selection [9,10]. In summary, FOXP2’s multidomain architecture - comprising structured DNA-binding and protein-interaction domains interspersed with flexible regulatory regions - enables a complex interplay of molecular interactions. Deciphering how these pieces come together is crucial for understanding FOXP2’s function as a transcriptional regulator.

Full-length FOXP2 has evaded structural resolution via traditional methods, likely due to its disorder and oligomeric flexibility.. Likewise, structural information on FOXP2 in complexes with DNA beyond the isolated FHD, or in complexes with other proteins, has been scarce. Biochemical studies have inferred some aspects: for instance, FOXP2’s LZ is required not only for dimerization but also for high-affinity DNA binding [10], and FOXP2 can heterodimerize with its paralogs FOXP1 and FOXP4. FOXP2 (and FOXP1) also physically interact with chromatin modifiers and other transcription factors in neurodevelopmental gene networks [3]. Furthermore, emerging evidence suggests that FOXP2 may form complexes with a variety of partners, including FOXP1/4, cell-cycle regulators like CDK3 (with its partner cyclin CCNC), and transcriptional co-repressors such as CtBP1/2 and TLE1/5. These interactions imply that FOXP2 cooperates with other developmental regulators to control gene expression programs. Consistent with this, pathogenic mutations in FOXP2 (such as R553H) can disrupt protein-protein interactions [4] or disturb the DNA-binding [11]. Overall, FOXP2 functions at the nexus of multiple molecular interactions - DNA binding, homomeric/heteromeric protein assemblies, and cofactor recruitment - making it a prime candidate for an integrated structural analysis.

Recent breakthroughs in deep learning-based structure prediction—exemplified by RoseTTAFold2 [12] and AlphaFold [13]—have transformed structural biology by enabling high-accuracy, proteome-scale modeling. As a result, nearly every protein sequence now comes with a plausible 3D structural model [14]. Notably, the latest iteration, AlphaFold3, extends these capabilities to model complex biomolecular interactions, including protein-DNA, protein-RNA, and multimeric protein assemblies, with remarkable accuracy [15]. AlphaFold3 is a next-generation deep learning model that substantially improves upon AlphaFold2’s protein structure predictions by incorporating a diffusion-based architecture and explicitly modeling multicomponent complexes [15,16]. These developments were recognized with the 2024 Nobel Prize in Chemistry.

Here, we apply AlphaFold3-multimer to systematically map FOXP2’s structure and interaction landscape. We predicted its full-length and oligomeric structures, modeled FOXP2-DNA binding with diverse sequences, and explored its interactions with known and candidate protein partners. Confidence was assessed via a scoring scheme based on residue-residue contact reproducibility across 25 AlphaFold3 replicates per complex. We integrated these models with disorder predictions, disease variants, DNA preferences, and known interactions to: (1) elucidate full-length FOXP2 architecture and oligomerization, (2) characterize its DNA binding with sequence specificity, (3) evaluate structural impacts of R553H and human-specific substitutions, and (4) model its protein interaction network. This integrated analysis highlights how FOXP2 functions via cooperative DNA recognition, multimerization, and selective cofactor recruitment, providing a framework for future experimental validation in neurodevelopment and evolution.

## Results

### Full-length FOXP2 structure

Using AlphaFold3, we predicted the structure of the full-length human FOXP2 protein (715 amino acids). Figure 1A and Figure 2A show the domain organization of FOXP2 from N- to C-terminus. The N-terminal segment (residues ∼1-100) contains the polyglutamine tract and is largely disordered in the monomer model. Following this, residues ∼101-250 form two long α-helices connected by a short turn; interestingly, this helical region corresponds to a portion of the so-called N-terminal repression domain and includes several known FOXP2 nonsense variants associated with childhood apraxia of speech (e.g., Q187*, Q196*, and Q210*). Residues 250-260 form a short linker leading into a central helix (residues ∼261-270) followed by a large unstructured loop (270-345) that contains the two human-specific substitutions (N303T and N325S,[1, 18]).

**Figure 1.**
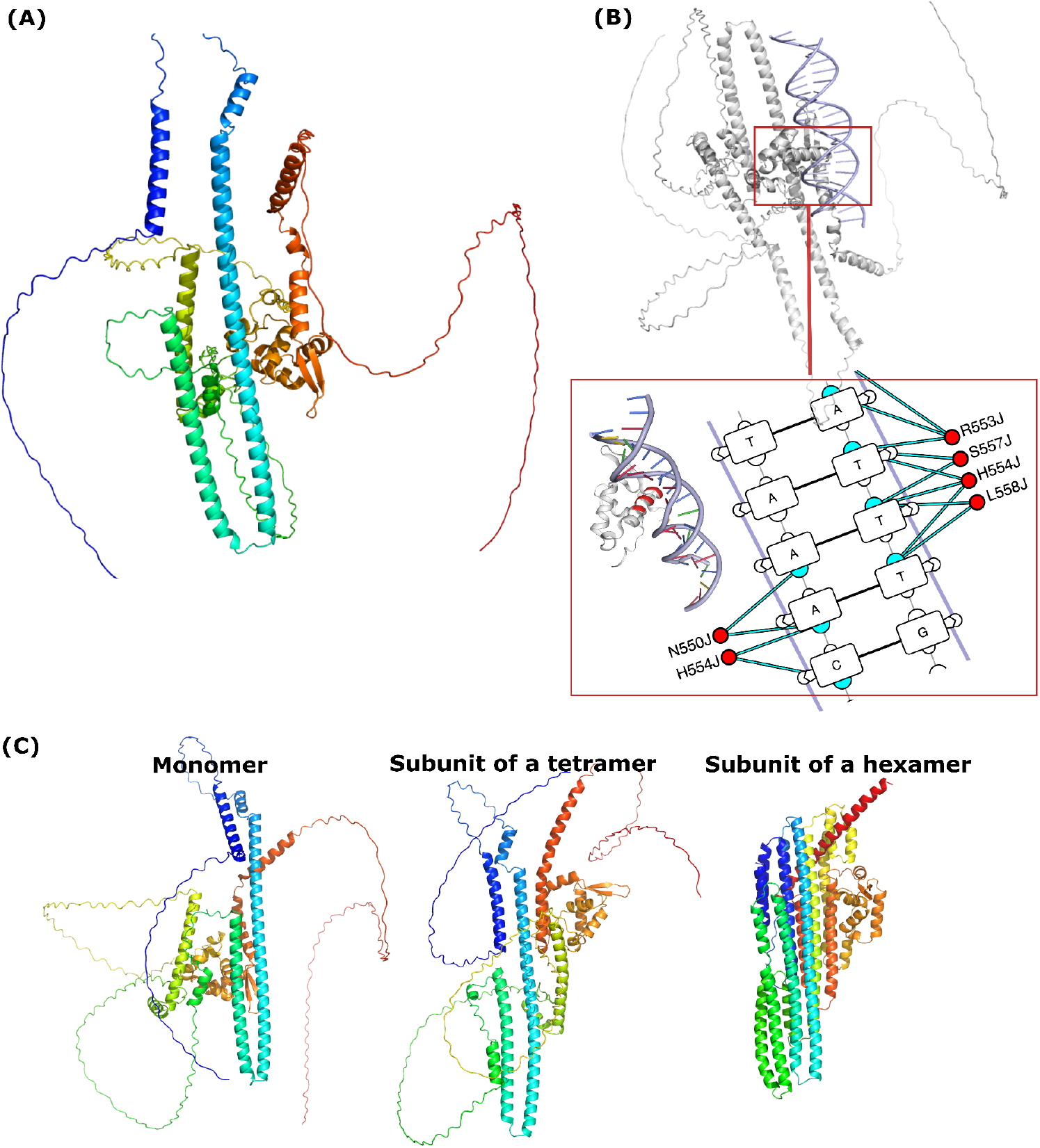
Full-length FOXP2 AlphaFold3 model and oligomerization. **A)** Monomeric AlphaFold3 model of FOXP2, colored by residue position from blue at the N-terminus to red at the C-terminus, illustrating a spectral gradient across the full-length protein. **B)** Visualization of the FOXP2 forkhead domain (FHD) bound to DNA using DNAproDB [17], highlighting key protein-DNA interactions at the binding interface. **C)** Comparison of FOXP2 in different oligomeric states (AlphaFold3 models): monomer (left), subunit of a tetramer (center), and subunit of a hexamer (right), each colored by residue position from blue (N-terminus) to red (C-terminus).

**Figure 2.**
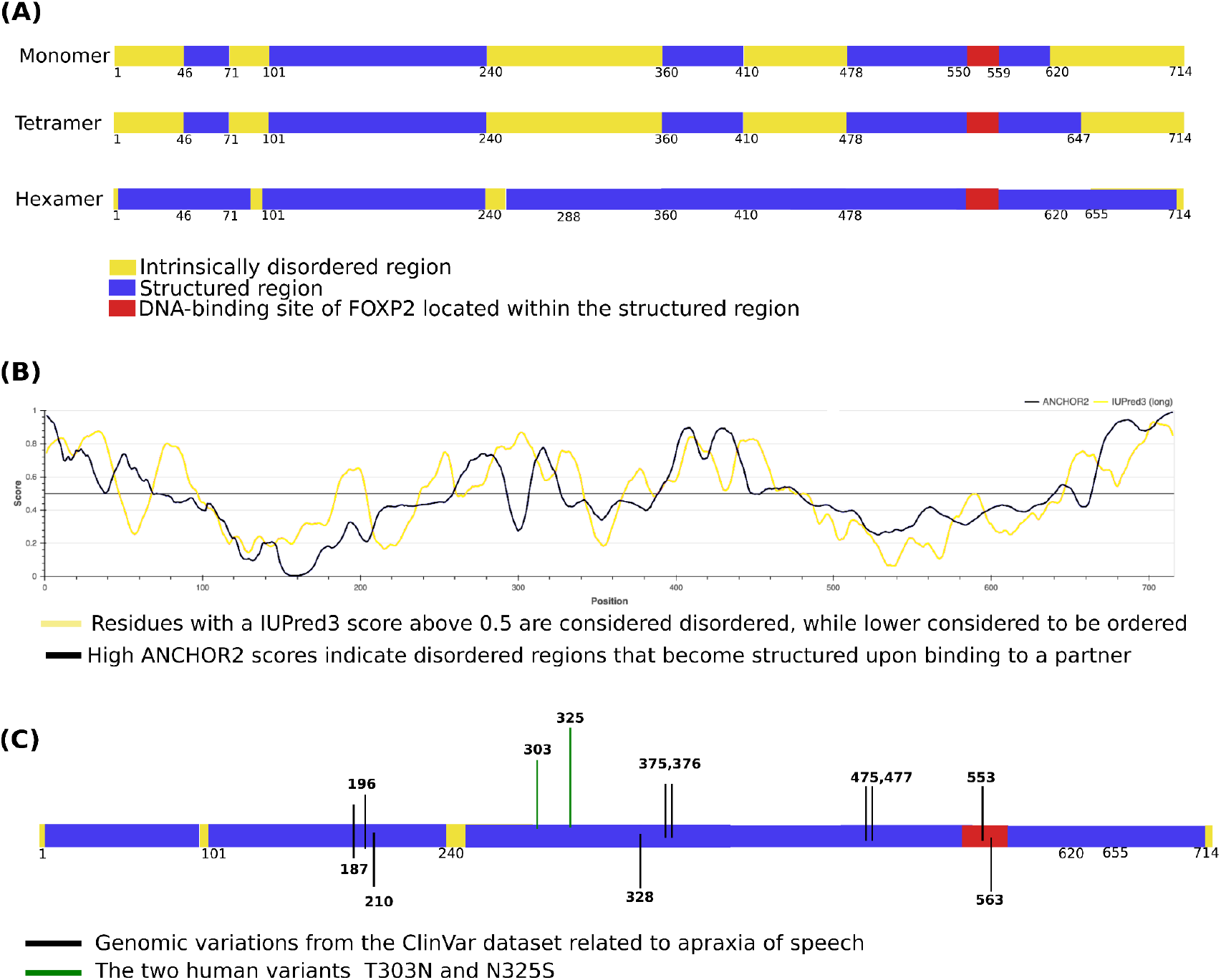
Structural and functional annotation of FOXP2. **A)** Schematic summary of FOXP2 structure in monomeric, tetrameric, and hexameric forms. Ordered regions are shown in blue, predicted intrinsically disordered regions in yellow, and DNA-binding sites in red. Oligomerization progressively orders regions predicted to be disordered in the monomer. **B)** Disorder prediction profiles for FOXP2. The yellow curve represents IUPred3 scores, where high values (typically >0.5) indicate residues likely to be intrinsically disordered. The black curve shows ANCHOR2 predictions, where high values (>0.5) suggest disordered regions with potential to mediate transient interactions with protein or DNA partners. These regions are candidates for structural transitions upon binding or complex formation. **C)** Mapping of pathogenic mutations and human-specific substitutions onto the FOXP2 structure. Variants cluster within regions predicted to be structured and functionally relevant, particularly in domains that may become stabilized upon oligomerization.

Next, the model resolves the zinc-finger (ZF, residues ∼346-366) and leucine-zipper (LZ, residues ∼367-400) motifs, which pack together to form an extended helical bundle (an “L”-shaped structure). Following the LZ, a flexible loop (residues 401-504) connects to the globular FHD (residues ∼505-585). The FHD adopts a compact winged-helix fold consisting of three α-helices (helix H3 being the DNA-recognition helix), followed by two small β-strands (the “wings”) and a C-terminal α-helix (helix H4) [8]. Finally, the extreme C-terminus of FOXP2 (residues 586-715) is an acidic tail rich in glutamate/aspartate; in the monomeric model, this tail is largely unstructured. The AlphaFold3 per-residue confidence (pLDDT) is high (>75%) for the ZF, LZ, and FHD regions, moderate (50-70) for the helix-turn-helix region (101-250) and parts of the 270-345 loop, and low (<50) for the poly-Q tract and the acidic tail, consistent with those regions being intrinsically disordered in isolation.

Importantly, our model’s FHD aligns closely with the experimentally determined FOXP2 FHD-DNA complex (PDB 2A07 [8]), with a root-mean-square deviation (RMSD) of ∼0.49 Å over 80 residues. The key DNA-contacting residues observed in the crystal structure (e.g., N543, R553, H554, S557, L558 in FOXP2 numbering) are correctly positioned in our full-length model’s FHD (these residues are highlighted in Figure 1B). This agreement increases confidence that the AlphaFold3 model has accurately captured the FOXP2 FHD conformation relevant for DNA binding.

### Oligomerization is predicted to induce ordered conformations in disordered regions

FOXP2 is known to form homodimers and also form higher-order oligomers in vitro [4,9]. We used AlphaFold3’s multimer to predict the structure of an FOXP2 dimer, tetramer, and hexamer (stoichiometries of 2, 4, or 6). The AlphaFold3 prediction for the FOXP2 hexamer converged on a well-defined homo-hexameric assembly (six FOXP2 monomers). In contrast, tetramer models showed minimal structural differences from the monomer (suggesting that a dimeric or dimer-of-dimers state does not fully engage the disordered regions) (Figure 2A). We therefore focus on the hexameric model. The FOXP2 hexamer is arranged as a trimer of dimers in a ring-like configuration (Supplementary Figure 1).

Within each dimer, the two FOXP2 protomers interact primarily via their LZ domains, forming the expected coiled-coil dimer interface (AlphaFold3 predicts a near-perfect LZ dimer in >90% of runs, consistent with the LZ being essential for dimerization [19]). In the hexamer, these three dimers further assemble such that their FHDs cluster toward the center of the complex, and the N-terminal helices and ZF-LZ regions extend radially outward. Oligomerization markedly orders regions that are disordered in the monomer.

In the hexamer model, the N-terminal segments (residues 50-80 of each protomer, immediately downstream of the poly-Q tract) from each pair of dimerized FOXP2 molecules form a long antiparallel coiled-coil with one another (Figure 1C). These N-terminal helices intertwine along the outer ring surface, stabilizing each dimer beyond the LZ contacts. Similarly, the large loop connecting the LZ to the FHD (residues ∼410-504) becomes ordered upon hexamerization: in the hexamer model, this loop in each protomer folds into a helical hairpin (two short helices around residues 420-480) that docks against the FHD of the neighboring protomer in the dimer. In other words, each FOXP2 protomer contributes a helical region that embraces the FHD of its dimer partner, thereby stabilizing the dimer interface in a DNA-dependent manner (see below). The C-terminal acidic tails (residues 586-715), which were largely unstructured in monomeric models, also adopt a more defined conformation in the hexamer. In the multimeric prediction, each acidic tail extends along the surface of its own protomer’s FHD and LZ, making multiple contacts (largely electrostatic) between acidic tail residues and basic patches on the FHD/LZ. Thus, in the context of the hexamer, the FOXP2 acidic tail appears to act in an intra-subunit autoinhibitory manner, consistent with prior proposals [9]. Overall, hexamer formation promotes a more compact, globular assembly of FOXP2. Previously flexible regions - the N-termini, the ZF-LZ linker, and the C-terminal tail - are reorganized into structured elements that participate in inter- or intra-subunit interfaces (Figure 1C and Figure 2A). This hexameric assembly is a computational prediction and has not yet been confirmed experimentally.

This mechanism of “coupled folding and assembly” is reminiscent of other IDR-containing proteins whose disordered segments become structured upon oligomerization or binding [19]. Our AlphaFold3 hexamer model is supported by experimental observations: analytical ultracentrifugation and mass photometry experiments have indicated that FOXP2 can form oligomers of ∼6-8 subunits in solution and increased helical content upon oligomerization [9]. Moreover, the ordered N- and C-terminal helices observed in the hexamer model align with regions of predicted disorder, suggesting that the model is capturing a biologically relevant, assembly-coupled folding event. The predicted hexamer provides a structural hypothesis for how FOXP2 may engage in cooperative DNA binding or chromatin looping, yet this remains to be tested experimentally.

Finally, to complement our structure-based modeling, we performed sequence-based predictions of intrinsic disorder using IUPred3 and ANCHOR2, a computational strategy we refer to here as ProbePLe. IUPred3 identifies intrinsically disordered regions (IDRs) based on energy estimation from amino acid sequences, while ANCHOR2 detects regions likely to undergo disorder-to-order transitions upon binding [20,21]. Both methods consistently predicted extensive disorder across the N-terminal and C-terminal tails and in the central region spanning residues ∼260-360, aligning closely with AlphaFold3’s monomeric model. Notably, ANCHOR2 highlighted some regions as likely to gain structure upon interaction, consistent with their reorganization into helices and interfaces in the hexameric form predicted by AlphaFold3 (Figure 2B). This convergence between sequence-based and structure-based approaches reinforces the interpretation that FOXP2 undergoes coupled folding and assembly during multimerization.

### DNA binding specificity mirrors experimental affinities

We next examined whether AlphaFold3 could predict FOXP2’s DNA-binding specificity across different target sequences. The FOXP2 FHD is known to recognize a DNA core motif with variable affinity depending on flanking sequences. Webb et al. (2017) used SPR to measure FOXP2 FHD binding kinetics across DNA sequences, proposing a three-tier model: low, moderate, and high affinity [22].

To reproduce and interpret these sequence preferences computationally, we used AlphaFold3 to model FOXP2-DNA complexes for multiple DNA sequences. We first validated the approach by modeling FOXP2 (monomer) bound to the 15 bp DNA from the FOXP2-DNA crystal (PDB 2A07) [8]. AlphaFold3 accurately reproduced the FHD-DNA geometry and key residue-base contacts (e.g., R553 to guanine; Figure 2B).. Encouraged by this, we modeled FOXP2 in a complex with five different 21-23 bp DNA sequences that had been characterized by Webb *et al*. [22]. These included two high-affinity (“Nelson”, “Zhu”; K_d ∼10-11 nM), two moderate (“Webb”, “Enard”; ∼17-19 nM), and one low-affinity (“Wang”; ∼55 nM) sequence. Sequence names reflect author surnames from original reports: “Nelson” [23], “Zhu” [24], “Wang” [25], “Enard” [26], and “Webb” [22]. We also included five control sequences of similar length with random bases (not expected to bind FOXP2 specifically).

AlphaFold3 produced plausible FOXP2-DNA complex models for all sequences. We computed FHD-DNA contact maps (averaged across 25 runs per sequence) to quantify the interaction footprint. The predicted DNA-binding ranking mirrored the experimental affinities (Table 1). The two high-affinity sequences (Nelson and Zhu) yielded models with dense and extensive contacts across the FHD dimer interface, particularly involving helix H3 residues R553, H554, and S557 in each protomer, making symmetric contacts with the DNA major grooves. The moderate-affinity sequences (Webb and Enard) showed intermediate contact density and slightly looser interfaces, consistent with their lower binding affinity. In contrast to the findings of Webb et al. [22], the low-affinity “Wang” sequence—despite exhibiting the weakest experimental binding—was predicted by AlphaFold3 to form a relatively extensive interface with FOXP2, ranking third in contact density. This discrepancy raises two possibilities: AlphaFold3 may capture a transient encounter complex that appears stable in silico but rapidly dissociates in reality due to fast off-rates, or alternatively, the “Wang” sequence may bind FOXP2 more effectively under specific experimental conditions not captured in the original SPR assay. *All five random sequences produced minimal contacts and low interaction probabilities*.

**Table 1.**
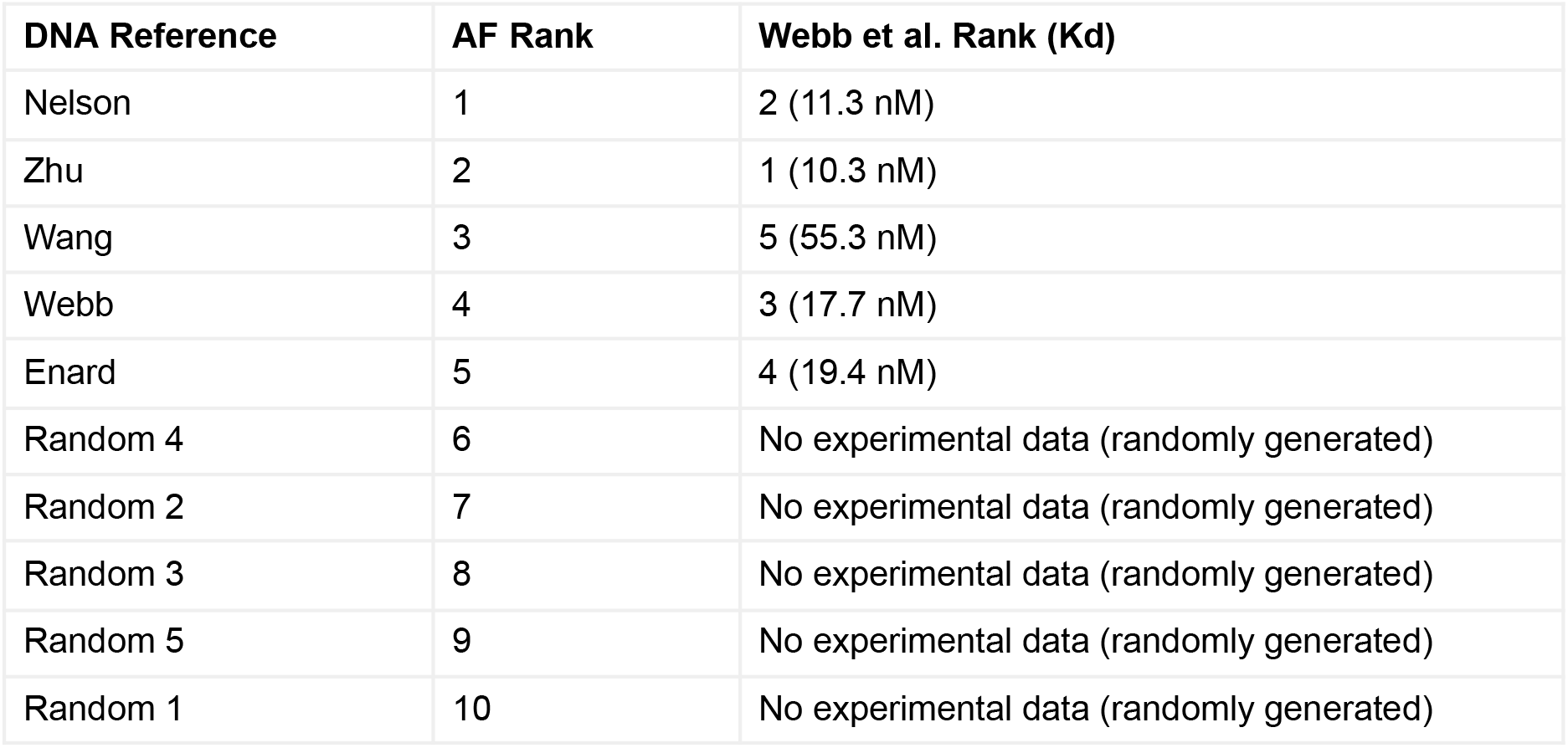
Comparison of AlphaFold-predicted FOXP2-DNA binding with experimental data. AlphaFold ranks represent predicted binding strength based on contact probability (1 = strongest). Experimental dissociation constants (Kd) were obtained from surface plasmon resonance (SPR) measurements reported by Webb et al. Predicted rankings show strong concordance with experimental affinity data for high-confidence DNA targets, except the Wang sequence, which showed low affinity experimentally despite a high predicted contact probability. Randomly generated sequences ranked lowest and lacked experimental binding data, supporting the specificity of FOXP2 for its consensus motifs.

| DNA Reference | AF Rank | Webb et al. Rank (Kd) |
| --- | --- | --- |
| Nelson | 1 | 2 (11.3 nM) |
| Zhu | 2 | 1 (10.3 nM) |
| Wang | 3 | 5 (55.3 nM) |
| Webb | 4 | 3 (17.7 nM) |
| Enard | 5 | 4 (19.4 nM) |
| Random 4 | 6 | No experimental data (randomly generated) |
| Random 2 | 7 | No experimental data (randomly generated) |
| Random 3 | 8 | No experimental data (randomly generated) |
| Random 5 | 9 | No experimental data (randomly generated) |
| Random 1 | 10 | No experimental data (randomly generated) |

We summarized contact-based rankings vs. SPR K_d values in Table 1. The AlphaFold3 predictions recapitulate the known specificity of FOXP2’s FHD for its cognate sites: sequences previously identified as high-affinity yield the most stable predicted complexes, whereas noncognate sequences do not induce stable binding. These align with the three-tier model of Webb et al. [22], in which FOXP2 can scan DNA in a partially disordered state and only locks into a fully folded dimeric complex on encountering optimal sequences.

### Structural impact of disease mutations

We next use our FOXP2 structural model to interpret known FOXP2 variants. Several FOXP2 mutations have been identified in individuals with speech and cognitive impairments [27,28]. We mapped clinically significant FOXP2 variants indexed in ClinVar [29] onto our structural models (Figure 2C). Such structural mapping offers testable hypotheses for how specific mutations might alter FOXP2 function (e.g., by destabilizing the hexamer, or by disrupting DNA contact, etc.). For example, the R553H missense mutation in the FHD is recurrently associated with speech apraxia [6]. In our model, R553 is located at the protein-DNA interface: it is one of the critical arginine residues on helix H3 that inserts into the DNA major groove [8]. Substituting R553 with histidine would remove the guanine-contacting side chain and likely disrupt one of the key electrostatic interactions anchoring FOXP2 to DNA. Indeed, our AlphaFold3 DNA-binding simulations for an R553H mutant predict markedly reduced DNA contacts and a loss of the tight FHD-DNA interface, consistent with a loss-of-function effect on DNA binding (Table 2).

**Table 2.** Contact probabilities of FOXP2 wild-type and R553H mutant. Residue-residue contact probabilities were predicted using AlphaFold3 for FOXP2 modeled in complex with DNA. The R553H substitution leads to reduced contact probabilities at the mutation site and across neighboring residues, suggesting a broader destabilization of the DNA-binding interface.

| Residue Pair | Contact Probability (Wild-Type) | Contact Probability (R553H) |
| --- | --- | --- |
| A_550-B_8 | 0.78 | 0.61 |
| A_550-B_9 | 0.73 | 0.52 |
| A_551-B_7 | 0.69 | 0.41 |
| A_553-C_12 | 0.75 | 0.67 |
| A_553-C_13 | 0.86 | 0.69 |
| A_554-B_7 | 0.62 | 0.48 |
| A_554-B_8 | 0.70 | 0.61 |
| A_554-C_15 | 0.51 | 0.38 |
| A_557-C_13 | 0.73 | 0.67 |
| A_557-C_14 | 0.68 | 0.59 |
| A_557-C_15 | 0.57 | 0.39 |

We also mapped the locations of the two human-specific FOXP2 substitutions, T302N and N325S [2,5], which reside within the long helices of the N-terminal region (residues 101-250). In the monomeric model, these helices pack against each other intramolecularly. However, in the hexameric model, they participate in inter-subunit N-terminal coiled-coil interactions. Notably, both positions 302 and 325 are situated at the interface of this coiled-coil, raising the possibility that these human-specific changes influence FOXP2 oligomerization dynamics or stability.

### Stable interactions with FOXP1, FOXP4, Pin1, and CDK3 in a DNA-dependent manne

Beyond its self-assembly and DNA binding, FOXP2 is known to engage with a variety of proteins as part of its broader regulatory roles [30]. We used AlphaFold3 to model binary complexes between FOXP2 and 12 known or candidate interactors obtained from the IntAct database [31] through UniProt [32] (Supplementary Table 1). This set included its paralogs FOXP1 and FOXP4; the prolyl isomerase Pin1; the cell cycle kinase CDK3; the co-repressors CtBP1 and CtBP2; the chromatin regulators SATB2 and SOX5; the nuclear hormone receptor NR2F2; the E3 ubiquitin ligase LNX1; the transcriptional co-repressor TLE5; and FAM124A, a nuclear protein recently linked to cortical transcriptional networks involving FOXP2. For each FOXP2 partner pair, we employed AlphaFold3-multimer and performed 25 independent prediction runs, each initiated with a different random seed. This approach introduces stochastic variation in the model initialization and MSA sampling, allowing us to assess the consistency and robustness of the predicted interfaces across replicates.

Four of the 12 partners—FOXP1, FOXP4, Pin1, and CDK3—yielded high-confidence interactions, each showing dependence on FOXP2’s DNA-binding state. Table 3 summarizes the interaction strength and reliability (median interface contact probability) for each partner ± DNA. FOXP1 and FOXP4 *formed stable heterodimers with FOXP2 (median contact probability >0*.*9), independent of DNA*. The FOXP2-FOXP1 and FOXP2-FOXP4 interface closely resembled the FOXP2 homodimer interface, centered on the LZ coiled-coil (Supplementary Figure 2). AlphaFold3 predicts that FOXP1 and FOXP4 can heterodimerize with FOXP2 via a nearly identical LZ interaction as the FOXP2 homodimer, *consistent with prior data and shared FOXP domain architecture* [9,10]. In our models, FOXP1/4 heterodimers also include contacts between the N-terminal helices and the LZ of the partner, analogous to the FOXP2 homodimer. These interactions were stable in both +DNA and -DNA models, *though DNA modestly enhanced interface tightness, possibly by aligning the dimer*. The DNA independence of FOXP1/4 interactions is expected, since their dimerization is driven by the LZ, which is unaffected by DNA. In contrast, interactions of FOXP2 with Pin1 and CDK3 were strongly DNA-dependent.

**Table 3.** Predicted interaction strength (AlphaFold3 contact probability) for each partner with and without DNA. Confidence and reliability scores were derived from 25 independent AlphaFold3 multimer predictions for each FOXP2 complex, modeled with and without DNA. Confidence represents the median residue-residue contact probability across models. Reliability reflects the consistency of contact patterns across random seeds. DNA presence enhances interaction stability for select partners, while several candidates showed no interaction under either condition.

| Protein | DNA in the complex |  | No DNA |  | Comment |
| --- | --- | --- | --- | --- | --- |
|  | Probability | Reliability | Probability | Reliability |  |
| FOXP1 | 0.98 | 1.00 | 0.92 | 0.99 | Strong interaction, stable with or without DNA |
| FOXP4 | 0.98 | 1.00 | 0.86 | 1.00 | Structurally similar to the FOXP1 interaction |
| PIN1 | 0.74 | 0.82 | 0.55 | 0.89 | DNA enhances interaction stability |
| CDK3 | 0.56 | 0.72 | -- | -- | Interaction observed only in the presence of DNA |
| F124A | 0.68 | 0.16 | 0.66 | 0.61 | Moderate; low reproducibility |
| TLE5 | 0.62 | 0.09 | 0.62 | 0.30 | High probability, low consistency |
| TSACC | 0.75 | 0.20 | 0.63 | 0.22 | High probability, low consistency |
| LNX1 | 0.60 | 0.20 | -- | -- | Weak and variable binding |
| CTBP1 | -- | -- | -- | -- | No interaction detected |
| CTBP2 | -- | -- | -- | -- | No interaction detected |
| SDCB1 | -- | -- | -- | -- | No interaction detected |
| CCNC | -- | -- | -- | -- | No interaction detect |

Pin1 recognizes phosphorylated Ser-Pro or Thr-Pro motifs in its targets. AlphaFold3 predicted a moderate affinity interaction between Pin1 and FOXP2 that was significantly stabilized in the DNA-bound state of FOXP2. CDK3, a cyclin-dependent kinase, was predicted to bind FOXP2 only in the DNA-bound context. In the +DNA models, CDK3 snugly fits into a pocket formed between FOXP2’s FHD and LZ (a groove that is partly created by the ordered 420-480 helical region when FOXP2 is dimerized on DNA). CDK3 made contacts primarily with FOXP2 residues 494-506 (within the FHD-LZ linker region). The interaction had a moderate probability (∼0.56) but high consistency (reliability ∼0.72) when DNA was present. In DNA models, FOXP2 rarely interacted with CDK3, and no stable complex was predicted. This implies that FOXP2 likely recruits CDK3 only when FOXP2 is DNA-bound. FOXP2 may recruit CDK3 to target DNA loci, though this prediction remains to be experimentally validated. A similar DNA-dependent interaction trend was observed for FAM124A with different binding sites. FAM124A reproducibly interacted with FOXP2 within a distinct site spanning residues 260-269. All binding sites, contact probabilities, and reliability scores are in Supplementary Table 2.

For the remaining candidate interactors (CtBP1, CtBP2, SATB2, SOX5, NR2F2, LNX1, and TLE5), AlphaFold3 did not predict any consistently stable interaction with FOXP2. In some individual runs, putative interfaces were observed—for example, occasional models showed FOXP2-TLE5 contacts along the N-terminal region—but these interactions were not reproducible across different seeds, with low reliability scores due to high variability in the predicted interfaces across replicates, reflecting a lack of consistent residue-residue engagement. It is important to note that the absence of a confidently predicted complex does not necessarily imply that the interaction is biologically irrelevant or false. Rather, it may suggest a transient, weak, or context-dependent interaction that AlphaFold3, designed to capture stable, high-affinity complexes, is not well-suited to model with confidence. These interactions may also depend on additional cofactors, post-translational modifications, or involve short linear motifs within disordered regions, which structure predictors often fail to resolve. Overall, these interaction models are testable hypotheses that remain to be verified in experiments.

## Conclusion

Our AlphaFold3 models predict that full-length FOXP2 is a dynamic molecule whose conformation depends on its oligomeric state and DNA binding. In the monomeric form, FOXP2 contains extensive intrinsically disordered regions—especially in the N-terminal segment, ZF-LZ linker, and acidic tail—which likely confer flexibility for DNA scanning and partner recognition. Upon oligomerization into a hexamer, these disordered regions undergo coupled folding, producing a highly ordered complex where protomers mutually stabilize each other. This higher-order assembly may enable allosteric regulation by aligning functional domains (LZ, ZF, FHD) for cooperative DNA engagement. Our models suggest that the FOXP2 hexamer may bind multiple DNA elements or bridge long DNA stretches, potentially enhancing chromatin association. AlphaFold3 also recapitulated FOXP2’s differential DNA sequence preferences and predicted suboptimal binding states, such as asymmetric FHD engagement, that may underlie fast dissociation at weaker sites. Mapping disease-associated variants revealed clustering at key interfaces, with the R553H mutation disrupting a critical DNA contact. Finally, our protein-protein interaction analysis identified a subset of FOXP2 partners that bind robustly in a DNA-dependent manner. Together, these findings integrate structural predictions with biochemical and evolutionary data, offering a comprehensive model of FOXP2 as a modular, multimeric transcriptional regulator. This work is a computational prediction and has not yet been confirmed experimentally.

## Methods

### AlphaFold3 structural modeling of FOXP2 and oligomers

Full-length human FOXP2 (UniProt Q9HCM9; 715 aa) was modeled using AlphaFold3 (v3.0.1). Predictions included the monomeric form as well as dimeric, tetrameric, and hexameric assemblies. Each oligomeric state was modeled in 25 independent runs using distinct random seeds to capture conformational variability and assess structural robustness. Structural metrics such as pLDDT (per-residue confidence) and predicted TM-score (pTM) were used for quality assessment. Interface formation and ordering of intrinsically disordered regions upon oligomerization were analyzed across models.

### Prediction of intrinsic disorder and binding-propensity regions

Intrinsic disorder in FOXP2 was evaluated using *IUPred3* and *ANCHOR2 [21]*. IUPred3 identifies disordered regions based on amino acid composition and estimated pairwise interaction energy, while ANCHOR2 predicts regions that are likely disordered but become ordered upon interaction. These predictions were compared to structural order in AlphaFold3 models to infer regions undergoing coupled folding upon binding or assembly. The service was accessed via the web server (https://iupred3.elte.hu).

### FOXP2-DNA interaction modeling

FOXP2-DNA binding specificity was analyzed by modeling FOXP2 monomers or dimers with 10 DNA sequences (five experimentally characterized targets and five GC-matched random controls). Each complex was modeled using AlphaFold3’s protein-nucleic acid mode. DNA was represented as double-stranded 21-23 bp polymers. Contact maps between FOXP2 residues (particularly in the FHD) and DNA bases were computed and averaged across replicates. Contact densities were used to derive binding affinity rankings, which were then compared to experimental dissociation constants (Kd) from SPR assays [22]. All analysis was performed in Python 3.

### Protein-protein interaction modeling

Twelve known or candidate FOXP2 interactors—sourced from the IntAct database—were modeled in binary complexes with FOXP2 using AlphaFold3-multimer. Each complex was simulated under two conditions: (1) DNA-free and (2) DNA-bound, using a known high-affinity 15 bp DNA sequence. For each condition, 25 replicate models were generated with different random seeds. Residue-residue contact probabilities and interfacial reproducibility across runs were used to assess interaction strength and confidence. High-confidence interactions were defined by consistent interfacial contact patterns in most of the runs with a median contact probability >0.5. All analysis was performed in Python 3.

### Mutational modeling

The pathogenic FOXP2 R553H variant was modeled in complex DNA to assess its effect on DNA-binding. Structural perturbations and loss of DNA contacts were quantified by comparing wild-type and mutant contact probabilities at the protein-DNA interface. All analysis was performed in Python 3.

### Visualization

Molecular visualizations were generated using PyMOL. DNA binding was visualised by DNAproDB [17].

## Supplementary Figures

**Figure 1.**
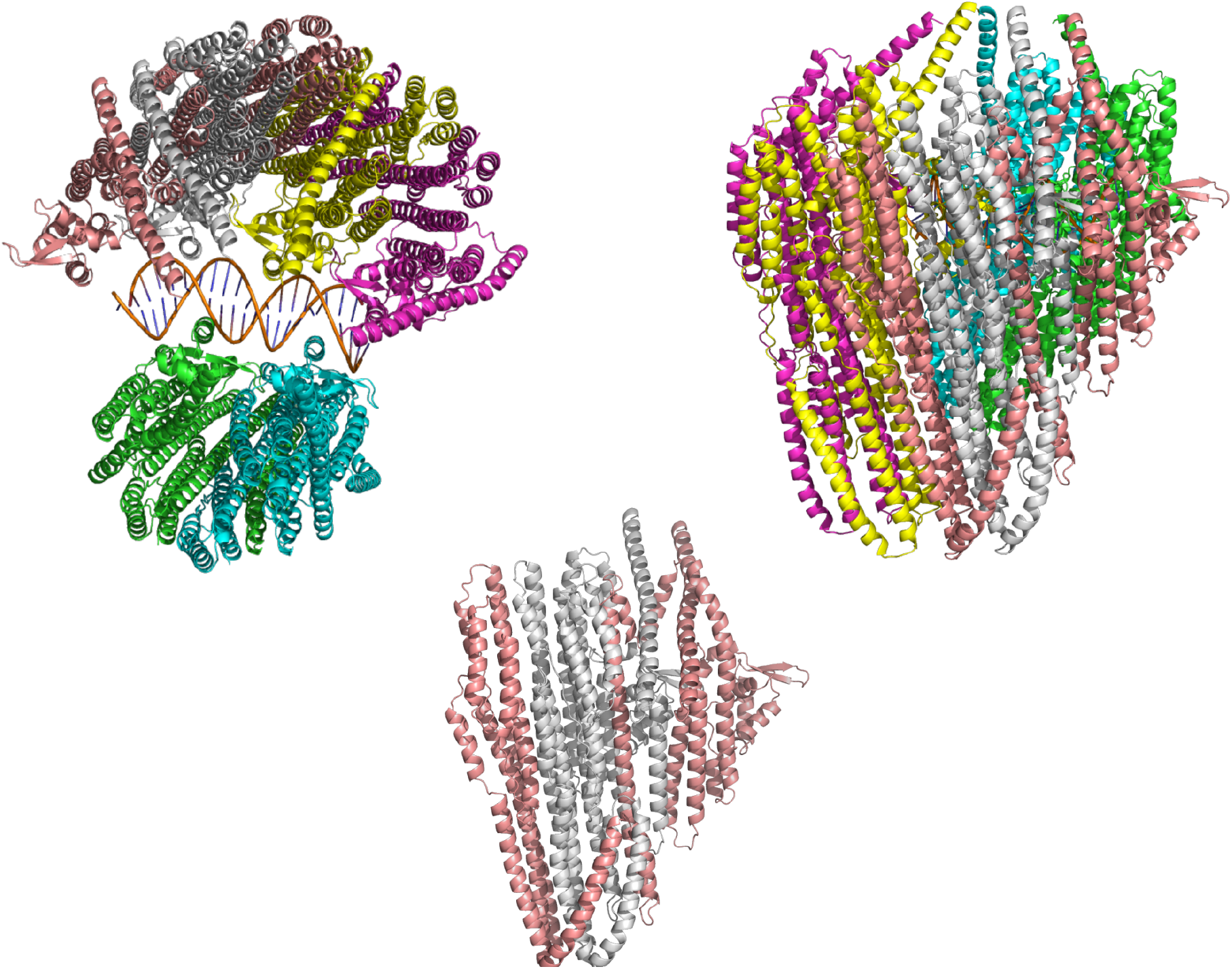
Predicted FOXP2 homo-hexamer structure. Top: Six FOXP2 protomers assemble into a ring-like hexameric complex, organized as a trimer of dimers. Each protomer is shown in a different color to highlight the overall symmetry and subunit arrangement. Bottom: One dimer from the hexamer was isolated to enhance visualization of the dimerization interface.

**Figure 2.**
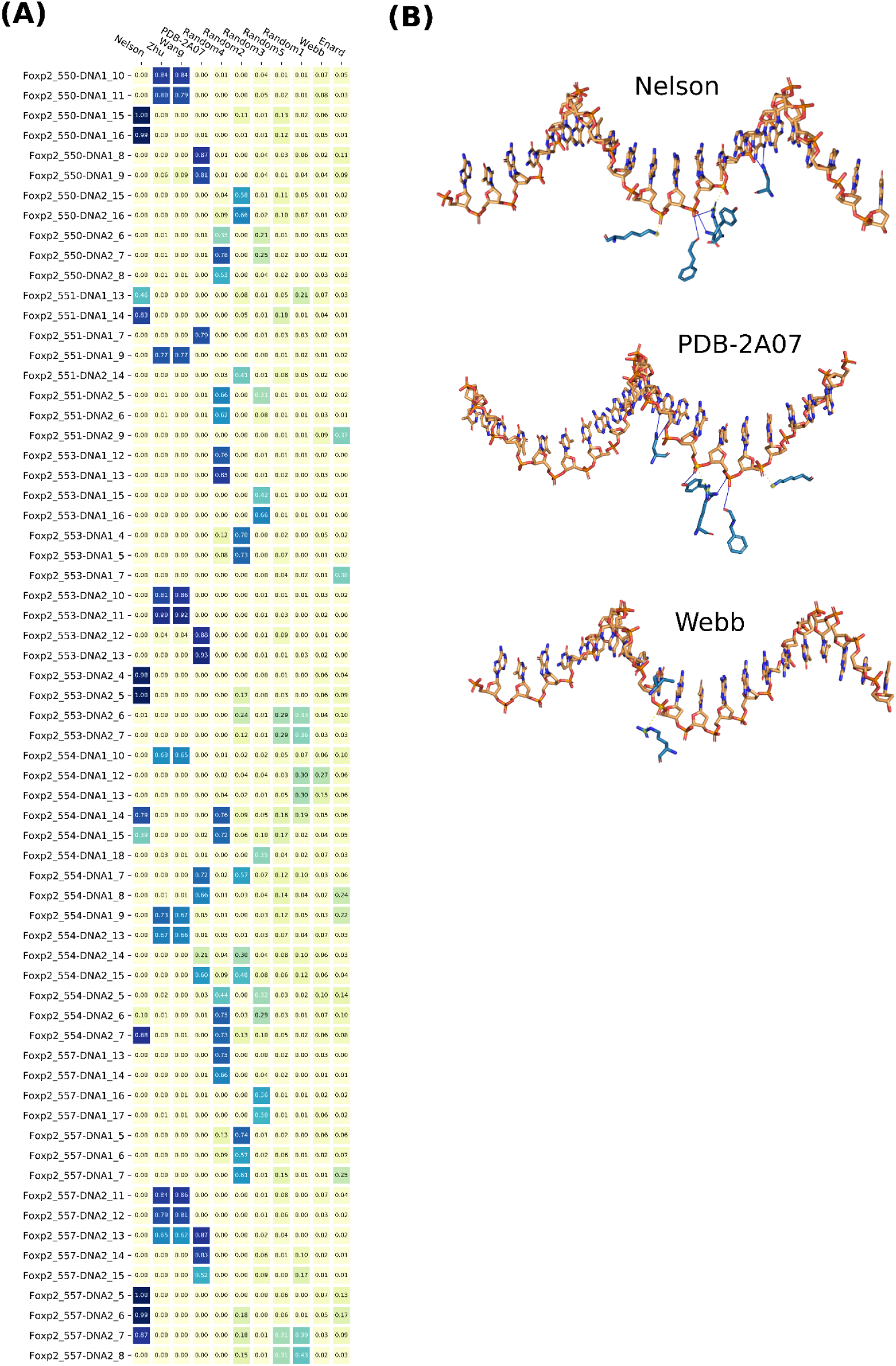
AlphaFold-predicted DNA binding specificity of FOXP2 compared with experimental data. A) Comparison of FOXP2-DNA binding predictions with surface plasmon resonance (SPR) measurements [22]. Left: heatmap showing aggregate contact probabilities at the FOXP2 forkhead domain (FHD)-DNA interface for five experimentally characterized DNA sequences (Nelson, Zhu, Webb, Enard, Wang, and a PDB 2A07 reference), alongside five randomly generated sequences (see Methods). B) Structural visualization of FOXP2 in complex with selected DNA sequences, illustrating key protein-DNA contacts.

**Figure 3.**
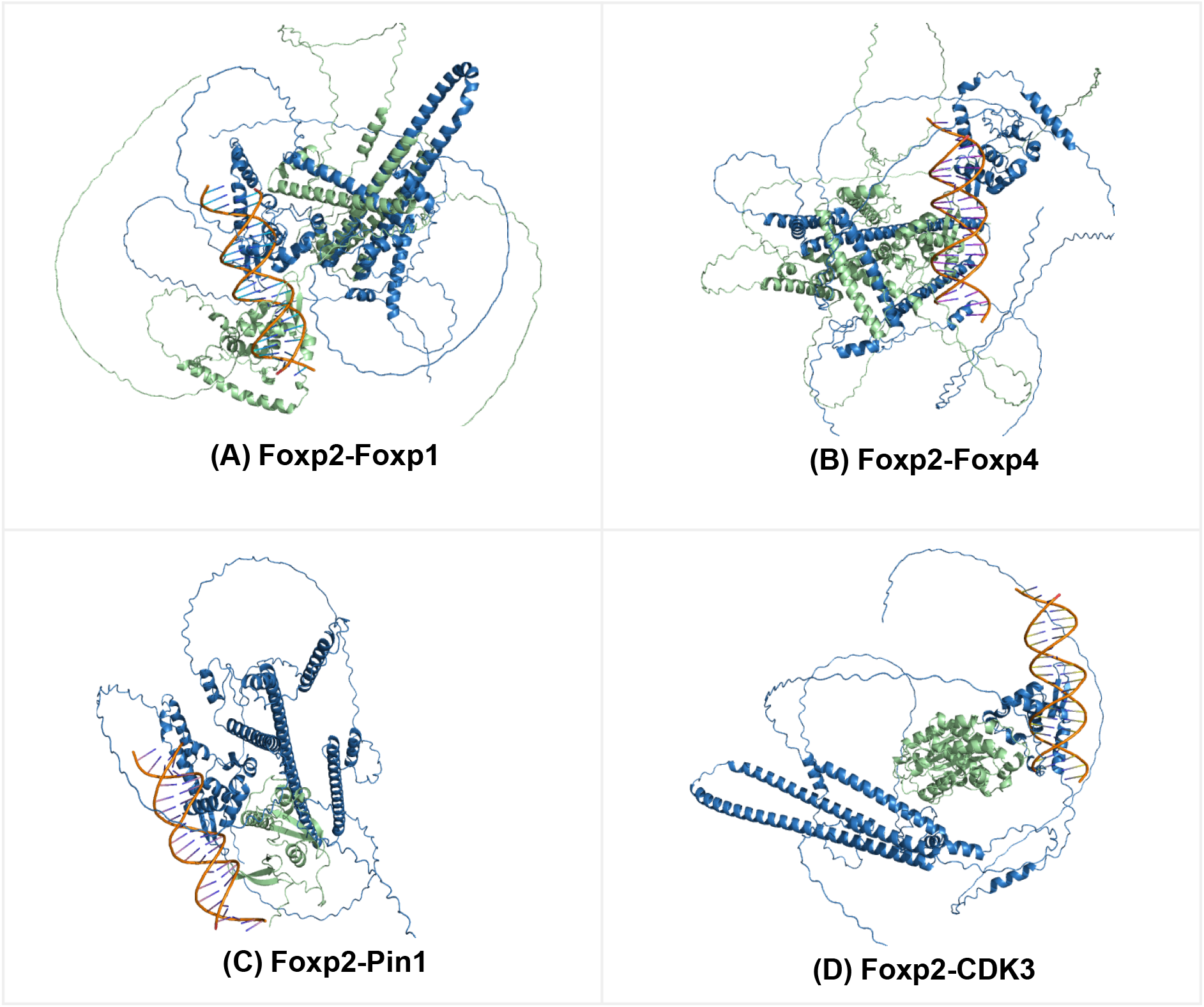
AlphaFold3-predicted protein-protein interactions involving FOXP2 in the presence of DNA.

## Supplementary Tables

**Supplementary Table 1:** Binary protein-protein interactions for 12 interactors identified from the IntAct database [31], accessed via UniProt.

https://sharing.biotec.tu-dresden.de/index.php/s/YMiRDFn3vF69PXB

**Supplementary Table 2:** FOXP2-protein interaction interface details. For each FOXP2 partner tested, this table lists: the top FOXP2 residues at the interface and binding sites

https://sharing.biotec.tu-dresden.de/index.php/s/FIDmkOk20G4UGJ7

